# Surgical hyoid bone repositioning effects on mandibular advancement and upper airway collapsibility

**DOI:** 10.1101/2024.09.12.612627

**Authors:** Hiba J Tannous, Corine J Samaha, Hassan A Chami, Joseph G Ghafari, Jason Amatoury

## Abstract

Mandibular advancement serves as a treatment option for obstructive sleep apnea (OSA), but its effectiveness differs among patients. The position of the hyoid bone is crucial for maintaining upper airway patency and may influence mandibular advancement outcomes. This study aimed to assess the impact of surgical hyoid re-positioning on mandibular advancement-induced changes in upper airway collapsibility in an animal model.

**Methods:** Twelve anesthetized male New Zealand White rabbits underwent mandibular advancement (0-4mm), combined with hyoid repositioning in various directions (anterior, cranial, caudal, anterior-cranial, anterior-caudal) and increments (0- 4mm). Upper airway collapsibility was quantified as the negative pressure required to close the airway (Pclose) at various mandibular and hyoid positions.

**Results:** Increasing mandibular advancement alone led to a progressive reduction in Pclose, indicating a decrease in upper airway collapsibility. Similarly, anterior hyoid repositioning alone resulted in incremental reductions in Pclose, with similar outcomes observed for anterior-cranial and anterior-caudal directions. When mandibular advancement was combined with anterior-based hyoid repositioning directions, a further decrease in Pclose was observed compared to when either intervention was applied alone. Cranial and caudal hyoid repositioning had no direct effect on Pclose or on mandibular advancement outcomes.

**Conclusions:** In summary, decreases in upper airway collapsibility induced by mandibular advancement are dependent on both hyoid repositioning direction and increment. The findings suggest that combining mandibular advancement with anterior-based hyoid repositioning may enhance the effectiveness of mandibular advancement in treating OSA.

## Introduction

The hyoid bone and mandible are crucial in maintaining the patency of the upper airway. Abnormal positioning of these bones can affect the mechanical behavior of upper airway tissues (including how they deform) and the effectiveness of pharyngeal muscles in responding to both static and dynamic upper airway loads (e.g. change in mandible position, intraluminal pressure variations, muscle activity) (1). Individuals with obstructive sleep (OSA) often exhibit an inferiorly positioned hyoid bone and a retruded mandible compared to healthy individuals (2–4). These anatomical deviations are associated with a more collapsible upper airway, a characteristic feature of OSA (3, 5, 6).

OSA is a highly prevalent disorder associated with serious health consequences, such as cardiovascular diseases and cognitive impairments (7–9). Accordingly, the treatment of OSA is a major health priority. Mandibular advancement, a treatment option for OSA that involves protruding the mandible using a dental appliance to keep the upper airway open during sleep, has been shown to reduce airway collapsibility (10). The success of mandibular advancement is reported in approximately 50% of patients, but the reasons for this variability are not well understood (11).

The mandible is a U-shaped bone forming the lower jaw, with a central body that supports the lower teeth and extends into the mandibular rami, which articulates with the skull at the temporomandibular joint (12). The hyoid bone, also U-shaped, is unique in that it does not articulate with any other bones. It has a central body and two pairs of processes: the lesser and greater horns. The hyoid is located at the base of the tongue, anterior to the epiglottis, and superior to the thyroid cartilage (13). While not directly involved in upper airway patency, the mandible is elevated and moved anteriorly by the lateral pterygoid, medial pterygoid, masseter and temporalis muscles. The mandible is connected to the hyoid bone via several pharyngeal muscles, including the genioglossus, geniohyoid, mylohyoid and digastric muscles that contribute to elevating and moving the hyoid anteriorly to open the airway (14, 15). Other muscles that connect to the hyoid include: the sternohyoid, thyrohyoid, and omohyoid muscles that pull the hyoid caudally; the middle constrictor and stylohyoid muscles that stabilize and move the hyoid posteriorly; and the hyoglossus that is involved in depressing and retracting the tongue (15, 16). As a result of the hyoid-mandible connections, a lower hyoid bone may decrease the effectiveness of mandibular advancement therapy due to alteration of muscle angles and/or altered tissue mechanical (stiffness) properties (1, 17). Hyoid repositioning surgeries, including those that raise the hyoid antero- cranially (hyomandibular suspension), are conducted to help compensate for the lower hyoid position in OSA (18). The combined effects of surgical hyoid bone repositioning and mandibular advancement in the treatment of OSA are unknown and require research to improve our understanding of this interaction and potentially enhance the treatment of OSA.

The aims of this study were to investigate the impact and interaction of hyoid bone repositioning and mandibular advancement on upper airway collapsibility using an anaesthetized rabbit model. The rabbit was chosen as an ideal model because of its fundamentally comparable upper airway to human anatomy whereby they share a freely suspended hyoid bone (19, 20), which differs from most non-primates in which the hyoid bone is relatively fixed (21).

## Material and methods

Studies were performed on 12 adult, male, New Zealand White rabbits weighing 2.9 ± 0.9 kg. The rabbits were bred and housed in the animal care facility at the American University of Beirut. The protocol was approved by the Institutional Animal Care and Use Committee (#19-08-544).

### Experimental setup

Most of the experimental methodology, except for that associated with mandibular advancement, has been previously reported (22). The experimental setup is illustrated in Fig 1.

**Fig 1.**
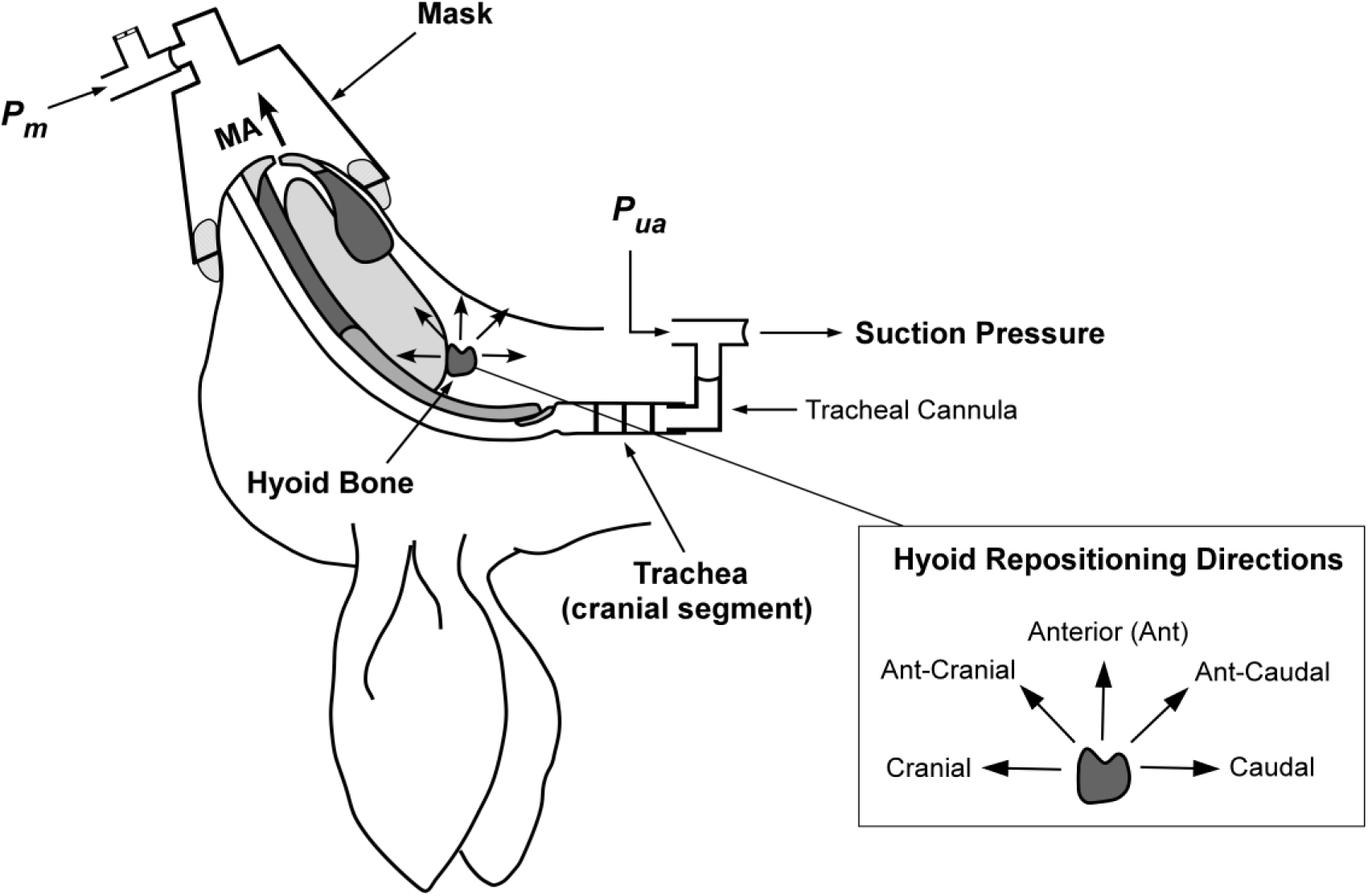
Schematic of the anaesthetized supine rabbit model. The upper airway is isolated at the level of the trachea. Mandibular advancement (MA) is applied in set increments. Hyoid displacement is applied to reposition the hyoid in the indicated directions (shown inset on right). Suction pressure (applied with a syringe) at the caudal tracheal end for closing pressure determination using upper airway pressure (Pua) and mask pressure (Pm). Figure adapted and modified from Samaha et al under the CC-BY license (22).

The rabbits were anesthetized with an intramuscular injection of ketamine (35 mg/kg) and xylazine (5mg/kg) followed by a continuous intravenous infusion of ketamine (15 mg/kg/hr) and xylazine (4.5 mg/kg/hr) to maintain anesthesia throughout the experiment. Heart and respiratory rates were monitored to ensure the rabbits’ physiological stability. At the conclusion of the experiment, the rabbits were euthanized using an anesthetic overdose.

The rabbits were positioned supine on a surgical platform. The head/neck position was controlled, such that a line drawn from the tragus to the external nares was at 50° to the horizontal. A ventral skin incision was made on the neck and blunt dissection was performed to expose the trachea. The baseline position of the trachea, taken between the fourth and fifth tracheal cartilage rings, was marked on the fixed experimental platform at the end of expiration.

The trachea was completely transected between the third and fourth tracheal rings to isolate the upper airway. This procedure resulted in the absence of airflow though the upper airway and the rabbits breathed spontaneously via the caudal trachea. An L-shaped tube was inserted into the caudal trachea and the pressure was monitored via this connection using a pressure transducer (Validyne DP45–32; Validyne Engineering, Northridge, CA). Another L-shaped tube was secured into the cranial trachea to reposition the tracheal segment to its pre-transection baseline position. A calibrated syringe and 100 cm volume extension for upper airway pressure application, and a pressure transducer (Validyne DP45–32) for measuring upper airway pressure (Pua), were also connected to the cranial L-shaped tube.

A small modified conical animal anesthetic mask (GaleMed VM-2, GaleMed, Taiwan) with inflatable sleeve was fitted to the rabbit’s snout to achieve a closed upper airway system. The mask allowed for a sealed system and the application of upper airway intra-luminal pressure and measurement of mask pressure (Pm) via a pressure transducer (Validyne DP45–32). To ensure a complete mask seal, the system was pressurized using a syringe. Pressure leaks were eliminated using petroleum jelly around the mask and inflation of the mask sleeve.

### Hyoid repositioning/displacement

A hyoid bone repositioning device was developed in-house to displace the hyoid bone in various increments and directions, as previously described (22). To attach the hyoid bone to the repositioning device, a miniscrew (RMO® Dual-Top, 2mm x 8mm) was inserted into the central part of the body of the hyoid. The screw served as a connection point between the hyoid bone and the clamp of the repositioning device, allowing for precise and controlled bone repositioning during the experiment.

### Mandibular advancement

A mandibular advancement splint (MAS) was custom made in-house based on plaster models of the rabbit’s maxilla (upper incisors) and mandible (lower incisors) (Fig 2). Alginate impression material was used to obtain a 3D impression of the models, which was then poured in white plaster. The MAS was constructed using cold-curing orthodontic acrylic resin (Vertex-Dental, AOPP2201000, shade 22, Netherlands) and incorporated an orthodontic expansion screw (Leone, A0890, Italy). The screw allowed for small gradual advancements of the mandible in the anterior direction.

**Fig 2.**
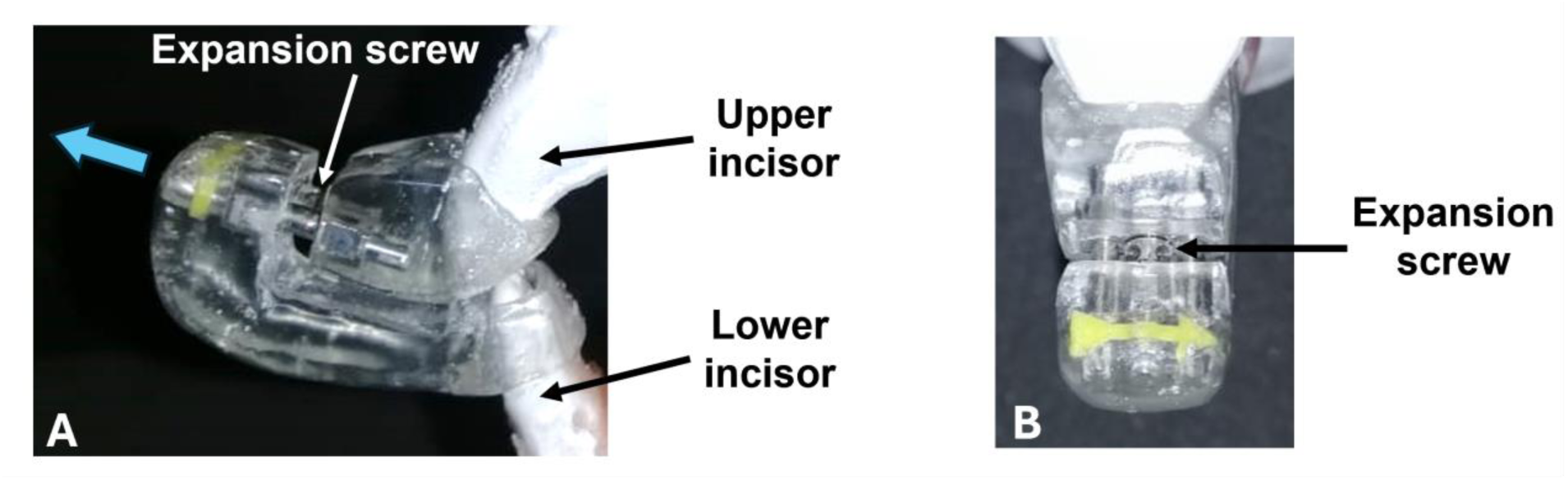
Mandibular advancement splint (MAS). (A) Side view of the MAS in which upper and lower incisors are inserted. The blue arrow indicates the direction of movement. The appliance is activated through an expansion screw that positions the lower incisor and mandible forward. (B) Frontal perspective of the appliance. The incorporated yellow arrow indicates the direction of screw activation.

The MAS was attached to the mandibular and maxillary incisors using glass ionomer luting cement (3M ESPE, self-curing, Germany). The MAS was fitted in such a way that the angle of mandibular advancement was 70° to the horizontal. By turning the screw in a clockwise direction, the mandible was displaced forward.

### Pclose measurements

The collapsibility of the upper airway was quantified using Pclose, which represents the closing pressure of the upper airway. When the upper airway was open, the pressure detected at the level of the mask (Pm) was equivalent to the pressure at the level of the trachea (Pua). Subsequently, a negative pressure was applied to the upper airway using the syringe connected to the cranial trachea. Pua and Pm were carefully monitored until the point of deviation, indicating the closure of the upper airway. The minimal pressure value reached by Pm before diverging from Pua was defined as Pclose (22). A more negative Pclose value indicates a less collapsible upper airway, while a less negative value indicates increased collapsibility.

### Interventional protocol

The hyoid bone was re-positioned within the mid-sagittal plane in sequence along anterior, caudal, cranial, anterior-cranial (45°) (ant-cranial), and anterior-caudal (45°) (ant-caudal) directions by 0, 2 and 4 mm. With each hyoid displacement, the mandible was advanced by 0, 2 and 4 mm. Pclose was measured for each hyoid repositioning direction/increment and mandibular advancement level. Following each Pclose measurement, the system was re-opened to atmosphere (0 cmH2O) and then closed again, ready for the next measurement.

### Data and statistical analysis

All physiological signals were acquired using a Power Lab 16 channel acquisition system and recorded using Lab Chart 8 (ADInstruments Ltd, Colorado, USA). The primary outcome, ΔPclose, was averaged for each rabbit for all 3 runs. Group averaged data were represented as mean ± SD. For mandibular advancement alone, the average of all runs before applying hyoid repositioning in each direction was included in the analysis. In combined mandibular advancement and hyoid displacement analysis, the mandibular advancement values prior to the hyoid repositioning intervention in a particular direction were considered baseline for direct relevance.

A fixed effects linear model (IBM SPSS v24) was used to analyze the effect of the three independent variables (hyoid repositioning direction and increment and mandibular advancement increment) on the outcome variable ΔPclose (dependent variable), as well as their interaction. A pairwise comparison was employed to assess which pairs of conditions significantly differed from one another. A Bonferroni post hoc test was performed to assess changes in key outcomes with hyoid and mandibular increments. Statistical significance for all the analyses was inferred for p < 0.05.

## Results

At baseline, Pclose was -4.2±0.4 cmH2O.When mandibular advancement was applied alone, Pclose was significantly decreased at each increment (p < 0.001; Fig 3). On average, Pclose decreased by -0.59 and -1.13 cmH2O at mandibular advancement levels of 2 and 4mm, respectively (Fig 3).

**Fig 3.**
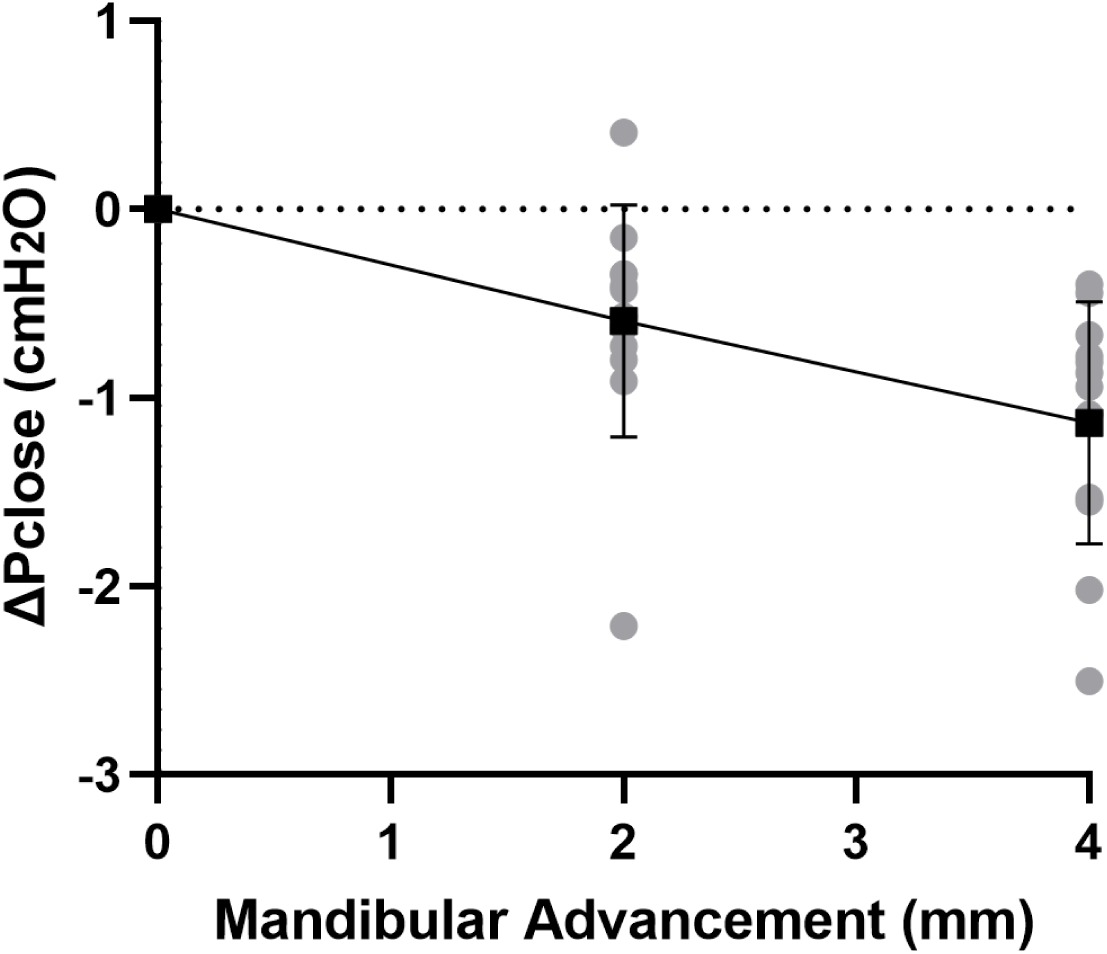
Change in closing pressure (ΔPclose) vs. mandibular advancement. Mandibular advancement alone caused a gradual decrease in ΔPclose with increasing increment. Individual rabbit (grey circles) and group mean values (black square) ± standard deviation (bars) are shown.

The changes in ΔPclose when both mandibular advancement and hyoid repositioning were combined are shown in Fig 5. There was no significant interaction between mandibular advancement and hyoid repositioning on Pclose in any direction (p>0.70). When mandibular advancement was combined with anterior, ant-cranial and ant-caudal hyoid repositioning directions, ΔPclose was more negative compared to when mandibular advancement was applied alone (Fig 5 A, D, E). For instance, a 4mm mandibular advancement combined with ant-cranial hyoid displacement resulted in a mean ΔPclose of -4.04 cmH2O compared with -1.43 cmH2O at 4mm mandibular advancement alone, and -2.82 cmH2O at 4mm ant-cranial hyoid displacement alone (Fig 5). No significant differences were observed between anterior, ant-cranial and ant- caudal hyoid displacement effects on mandibular advancement induced ΔPclose outcomes (p>1.0). Cranial and caudal hyoid displacement directions had no statistically significant effect on mandibular advancement (p>1.0).

Pclose showed progressive decrease with each increment in hyoid displacement in the anterior, ant-caudal and ant-cranial directions, reaching on average -2.31 to -2.82 cmH2O at 4 mm (p<0.001; Fig 4). The decrease in Pclose was not statistically significant between all anterior-based directions at corresponding increments (p>1.0). Pclose was not significantly altered when the hyoid was repositioned in cranial or caudal directions (p>1.0).

**Fig 4.**
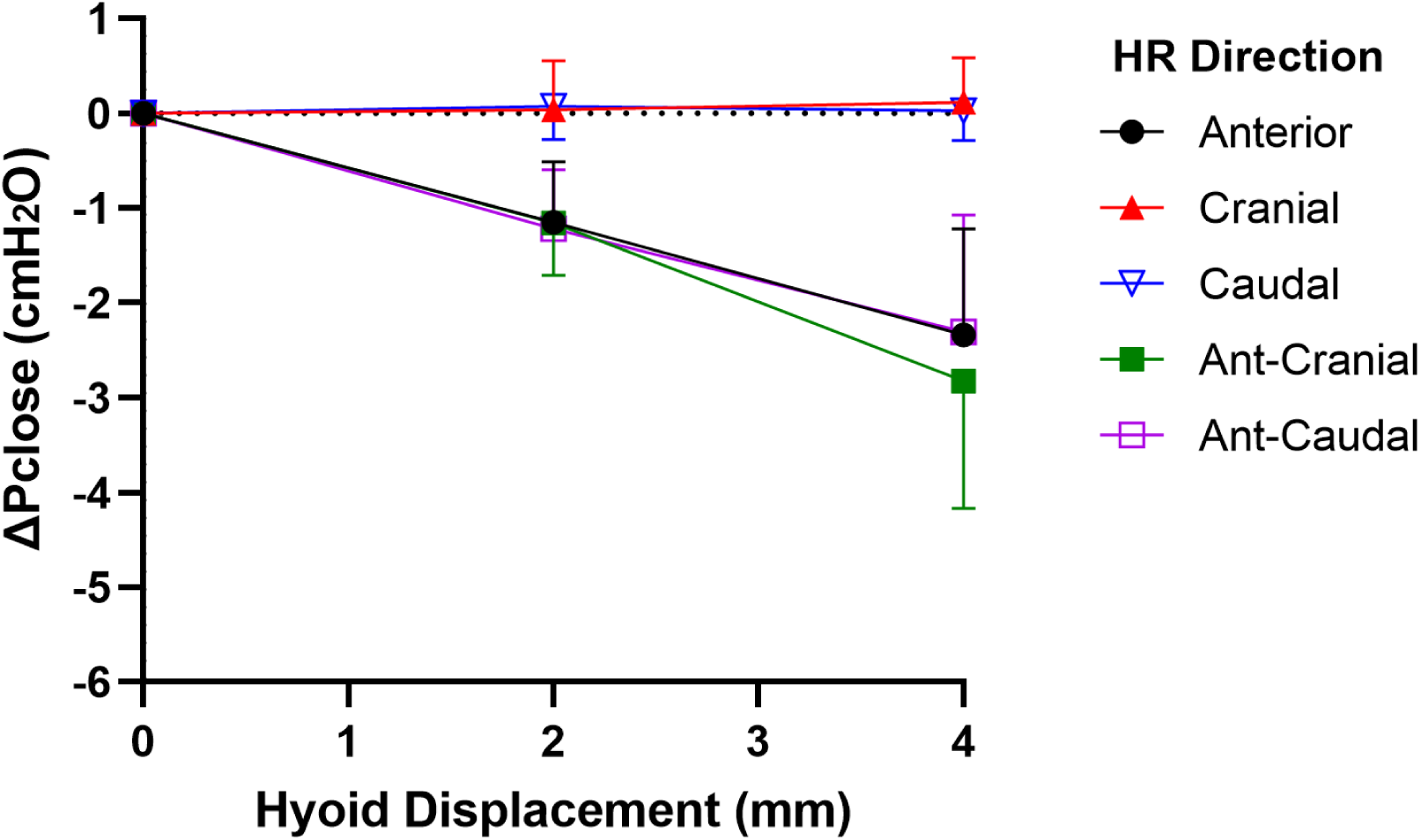
Change in closing pressure (ΔPclose) vs. hyoid repositioning (HR). Anterior (ant), ant- cranial and ant-caudal HR directions resulted in a gradual and similar decrease ΔPclose with each hyoid displacement. On the other hand, cranial and caudal HR directions had no significant effect on ΔPclose. Data are presented as mean group values (points) ± standard deviation (bars).

**Fig 5.**
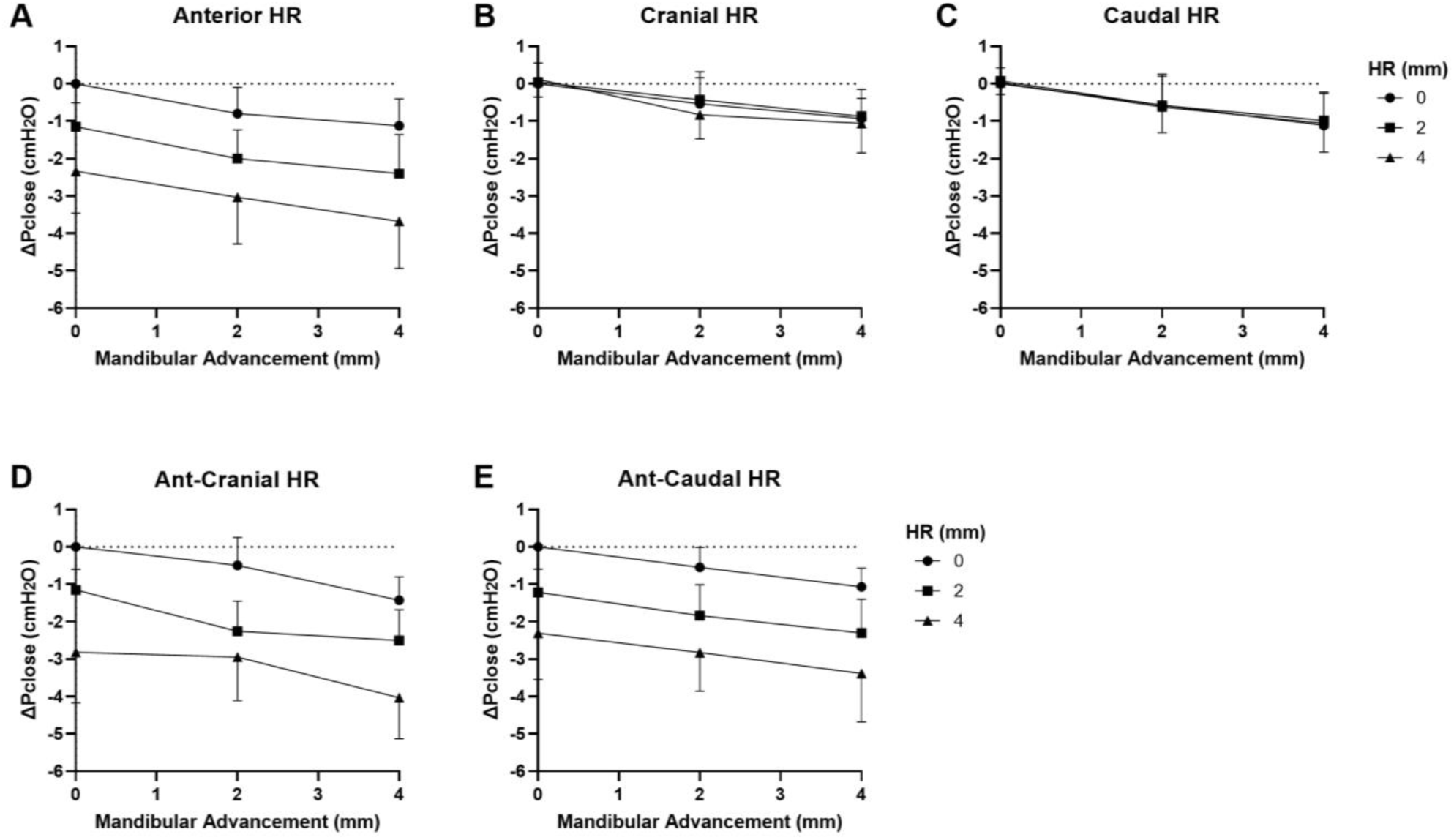
Change in closing pressure (ΔPclose) vs. mandibular advancement (MA) and hyoid repositioning (HR) combined. Increasing levels of MA at HR increments of 0, 2 and 4 mm are shown in A) anterior, B) cranial, C) caudal, D) ant-cranial, E) ant-caudal directions. When MA is combined with HR, there was an additive effect on ΔPclose such that it decreased even further then when either intervention was applied alone for anterior, ant-cranial and ant-caudal directions (A,D,E). However, ΔPclose did not significantly change with MA+HR for cranial and caudal directions compared to MA alone (B,C). Data are presented as mean group values (points) ± standard deviation (bars).

## Discussion

This study has demonstrated that both graded mandibular advancement and hyoid repositioning in anterior-based directions, but not in cranial or caudal directions, independently decreased Pclose and hence reduced collapsibility of the upper airway. When mandibular advancement was combined with anterior-based hyoid repositioning, the effect was additive, resulting in further decrease in Pclose than when mandibular advancement was applied alone. These outcomes suggest that the effectiveness of mandibular advancement in treating OSA may be improved when combined with hyoid bone repositioning in anterior-based directions.

### Mandibular advancement

Our findings are consistent with studies that showed reductions in upper airway collapsibility with mandibular advancement in humans (10, 23, 24). Studies have also demonstrated that mandibular advancement enlarges the upper airway (20, 25, 26) and can increase tissue stress/stiffness (20, 27), factors that could mediate the observed reduced collapsibility. Furthermore, mandibular advancement can alter the tongue’s dilatory movements to potentially improve therapeutic response (28). The effect of mandibular advancement on upper airway patency has also been partially related to the movement of the hyoid bone (20, 27, 29, 30).

When the mandible is advanced, the hyoid bone moves in an anterior/anterior-cranial direction (20, 30), which assists in redistributing the mandibular advancement load throughout the upper airway to enlarge and stiffen it (20, 27). Indeed, a computational finite element model of the rabbit upper airway showed that mandibular advancement effects on upper airway soft tissue displacements were reduced when the hyoid was fixed compared to when it was free to move (27). Nonetheless, even with the hyoid fixed in the current study, similar to surgical hyoid repositioning therapies for OSA, mandibular advancement can still reduce upper airway collapsibility. Mandibular advancement does not just impact the hyoid bone, but also alters the movement and stretch of the tongue (28) and other muscles and connections to the upper airway, like the pterygomandibular raphe (31), so that increases in upper airway patency and stiffening of soft tissues can occur. However, if the hyoid remained mobile when in its baseline position, it is possible that the effects of mandibular advancement would be greater (27).

### Hyoid repositioning

The independent effects of hyoid displacement on upper airway collapsibility observed in this study are consistent with our previous experimental results in rabbits (22). Anterior-based hyoid displacements progressively decreased upper airway collapsibility, while cranial and caudal hyoid displacements had no effect. Similar enhancements in upper airway patency through anterior and ant-caudal hyoid repositioning have also been observed in human and dog models (21, 32).

We previously hypothesized that the apparent lack of impact of cranial or caudal hyoid repositioning on upper airway collapsibility may be associated with compression/stretching of tissues above (e.g. genioglossus, geniohyoid, hyoglossus, styloglossus, stylohyoid and palatoglossus muscles) and below (e.g. thyrohyoid membrane and ligament, and thyrohyoid, sternohyoid and omohyoid muscles) the hyoid bone (22). It is likely that moving the hyoid cranially stretches upper airway tissues caudally, contributing to improved upper airway stability. However, the cranial hyoid movement compresses airway tissues cranially, leading to a more collapsible airway upstream. The opposite effect occurs with caudal repositioning. Consequently, the improvement in upper airway collapsibility in one segment is offset by a reduction in collapsibility in another, leading to an overall lack of change in collapsibility. In previous computational model simulations, we predicted that a decrease or lack of change in upper airway size with caudal/cranial hyoid repositioning likely contributes to the absence of change in Pclose observed with caudal/cranial hyoid repositioning experimentally (17) .

### Combined hyoid repositioning and mandibular advancement and clinical implications

To our knowledge, the combined effect of surgical hyoid repositioning and mandibular advancement has not been previously studied. The finding that combining hyoid repositioning in anterior-based directions with mandibular advancement leads to a more pronounced decrease in Pclose compared to either intervention alone is both novel and significant. For instance, the combination of mandibular advancement by 4mm and hyoid ant-cranial repositioning by 4mm yielded a 183% decrease in upper airway collapsibility compared to a 4mm mandibular advancement alone. This combination is particularly impactful as upper airway tissues and dilator muscles are pulled in approximately the same direction by both interventions. Thus, a treatment approach combining ant-cranial hyoid repositioning through hyomandibular advancement surgery, along with mandibular advancement, may improve airway patency in individuals who do not respond well to mandibular advancement alone. Additionally, combining surgical hyoid repositioning with mandibular advancement may reduce the excessive amount of mandible advancement required for effective OSA treatment, thereby potentially mitigating side effects such as temporomandibular or dental related pain, or dental/skeletal structural changes (33, 34). Further studies examining a combined hyoid repositioning and mandibular advancement therapy approach in humans are necessary to confirm these hypotheses.

Other clinical implications relate to issues not in the scope of this work and warrant separate investigation, such as the impact of hyoid bone and mandibular positions on swallowing function, as swallowing is a protective mechanism that prevents aspiration of saliva during sleep (35), and the effect on airway collapsibility of posterior mandibular orthognathic surgical movement.

### Critique of methods

General limitations of the current study with hyoid repositioning alone and other interventions have been detailed previously (19, 20, 22), and will be discussed briefly here.

#### Rabbit model

A rabbit model was used for the current study, which we and others have repeatedly adopted in investigating upper airway physiology and mechanics with demonstrated similarity of rabbit upper airway outcomes to the human circumstance (19, 20, 22, 27, 36–44). Although the rabbit’s craniofacial structure differs from that of humans and it possesses an overlapping soft palate and epiglottis, the general similarity of its upper airway structure makes it ideal for understanding concepts related to hyoid repositioning and mandibular advancement. Both interventions applied alone have shown comparable outcomes in the anesthetized rabbit to the sleeping or anesthetized human (20, 22, 27), which provides us with some confidence in the translatability of our combined intervention outcomes to the human Furthermore, a significant advantage of the rabbit model is its mobile hyoid bone, which lacks fixed bony attachments like the human, a characteristic not found in other non-primates such as dogs, felines and rats (19, 20). This allowed the hyoid to be readily repositioned in all directions and increments adopted in this study.

The natural position and morphology of the rabbit and human hyoid bones are very similar, with both situated directly below the tongue and possessing a central body with lesser and greater horn protrusions (13, 20, 45, 46). Furthermore, hyoid anatomical connections between both species are largely the same, except for two notable differences (16, 47): 1) In rabbits, the digastric muscle has a single belly that originates from the temporal bone and inserts directly onto the mandible, unlike in humans, where the muscle connects to the hyoid bone via a tendon. As a result, the hyoid is not influenced by the digastric muscle in rabbits as it is in humans. 2) The stylohyoid muscle in rabbits arises as two distinct heads, the stylohyoid major and minor muscles, which insert onto the greater and lesser hyoid horns, respectively. In humans, the stylohyoid muscle has a single head that connects to the grater horn of the hyoid bone, and the stylohyoid ligament in humans is replaced by the stylohyoid minor muscle in rabbits.

Given the primarily similar anatomical connections of the hyoid bone in rabbits and humans, it may be expected that the movement of the hyoid bone in both species is also similar under applied loads relevant to the upper airway. For instance, with mandibular advancement, the hyoid bone is displaced in a cranial-anterior direction in rabbits (19) and humans (30, 48). With increased lung volume/caudal tracheal displacement, the hyoid bone moves caudally in both species (19, 49). With activation of genioglossus and/or geniohyoid muscles, the hyoid moves anteriorly in both rabbits and humans (50, 51).

It is important to note that our rabbit model is not designed to replicate OSA but rather to simulate a healthy, well-controlled upper airway, which is one of the model’s major advantages to understanding fundamental mechanisms (see also below). OSA is a complex disorder influenced by multiple factors that require varied treatment approaches (52). We propose that, in certain individuals with OSA, this combination of mandibular advancement and anterior-based hyoid repositioning could be key to successful treatment. However, future studies in humans are needed to identify which individuals would benefit most from this approach.

An isolated upper airway preparation was used in this study with rabbits deeply anaesthetized, as per previous preparations (19, 20, 22). General anesthesia reduces upper airway muscle tone (53, 54). This reduction is ideal for our study, as general anesthesia creates an upper airway model similar to sleep, particularly in terms of collapsibility (54, 55). The ketamine and xylazine combination used in this study is extensively employed as an anesthetic agent in upper airway research (19, 20, 36, 56–58). Xylazine induces central muscle relaxation and anesthesia (59), while ketamine, in addition to providing sedation, counteracts any respiratory depression caused by xylazine, helping to maintain a relatively stable breathing pattern (60, 61). This combination is suitable for airway studies due to its ability to maintain near-passive airway characteristics, with upper airway muscle activity further reduced with the isolated upper airway in the current study, while preventing complete airway collapse (62, 63). The initial step in understanding the passive response of the upper airway to combined hyoid bone repositioning and mandibular advancement is necessary and advantageous for using animal models. By removing factors associated with upper airway muscle activity and airflow, we can first understand how hyoid repositioning and mandibular advancement impact the upper airway alone, and then comprehend in the future how muscle activity and upper airway airflow may alter outcomes in an intact upper airway (39, 64).

The reference point for the hyoid (hyoid repositioning = 0mm) and mandible (mandibular advancement = 0mm) represents the natural baseline position of these structures for each rabbit in the current setup (i.e., supine position, 50-degree head/neck angle, under anesthesia, etc.). The advantage of using a model based on healthy adult rabbits of the same species, gender, and similar age is that the hyoid and mandible are generally positioned similarly in each animal. However, we acknowledge that rabbits are not identical, and factors such as differences in body size, slight variations in positioning, or anesthetic level may have influenced the baseline reference positions of the hyoid and mandible, contributing to some variability in the study outcomes. Despite this, our primary focus was on investigating changes in Pclose with hyoid repositioning and mandibular advancement relative to each rabbit’s baseline position, rather than absolute values. This approach helps minimize noise associated with potential baseline position differences. Future studies incorporating imaging techniques to measure baseline hyoid and mandible positions could reduce potential variations in outcomes related to positioning between animals. Such imaging studies could not only be used to measure reference points, but also investigate changes in upper airway geometry and tissue morphology resulting from hyoid repositioning and mandibular advancement.

#### Hyoid bone setup

The hyoid bone was fixed in the new position and unable to move with any additional load, including mandibular advancement. This set-up is relatively similar to hyoid repositioning surgeries, in which the hyoid bone is fixed to the mandible or the thyroid cartilage. During normal functioning, the hyoid moves in response to various active and passive loads. Indeed, the hyoid bone is displaced with mandibular advancement (20, 30). Experimentally, preserving hyoid mobility after displacement in all directions investigated in this study is challenging. However, additional research, such as with computational modeling, could explore whether preserving hyoid mobility post-surgical repositioning may further enhance upper airway patency.

#### Statistics

We did not conduct a power analysis to determine the sample size for this exploratory animal study prior to its implementation. Instead, we used sample sizes from similar previous rabbit model investigations, both our own and those conducted by others, as a guide (19, 20, 22, 36, 58, 65). However, a post hoc power analysis revealed that for an alpha of 0.05, the power to detect differences in Pclose between increments of hyoid repositioning (for each direction) and mandibular advancement was at least 87%. The exception was the comparison between mandibular advancement of 2 mm and 4 mm, which had a power of 68%. Thus, the findings from this comparison should be interpreted with some caution.

## Conclusion

This study has demonstrated that combining mandibular advancement with anterior-based hyoid bone repositioning leads to further reduction in upper airway collapsibility compared to either intevention applied alone. However, no significant effect was observed with cranial or caudal hyoid repositioning. These findings suggest that indivudals with OSA who do not respond adequately to mandibular advancement alone may benefit from a combined therapeutic approach involving both mandibular advancement and anterior-based surgical hyoid repositioning. Such a combined treatment strategy holds promise for enhancing the management of OSA in these individuals. Further research and clinical studies are necessary to validate these findings in humans and refine OSA treatment protocols for personalized patient care.

## Acknowledgements

The authors would like to thank the staff of the MSFEA Mechanical Workshop for their assistance in experimental apparatus setup and Ms. Michella Samaan from the American University of Beirut Medical Center (AUBMC) Dental Laboratory for help with the development of the rabbit mandibular advancement splint. The authors would also like to thank Associate Professors Terence Amis and Kristina Kairaitis (The Westmead Institute for Medical Research and University of Sydney, Australia) and Professor Lynne Bilston (Neuroscience Research Australia and University of New South Wales, Australia) for their intellectual input and support.

## Notes

### Competing Interest Statement

The authors have declared no competing interest.

